# Cryptic Dispersal Networks Connect Habitat Patches in ways not Captured by Distance

**DOI:** 10.1101/341123

**Authors:** Rachel M. Germain, Natalie T. Jones, Tess N. Grainger

## Abstract

Species interact with the physical world in complex ways, and life history strategies might cause species to differ in how they experience connectedness of the same landscape. As a consequence, dispersal limitation might be present but not captured by distance-based measures of connectivity. To test these ideas, we surveyed plant communities that associate with serpentine soils but differ in dispersal mode (gravity, animal, or wind), and used satellite imagery to quantify forms of landscape connectivity associated with each dispersal mode. Our data yielded two key insights: First, dispersal limitation appeared to be absent using a conventional distance-based measure of connectivity, but emerged after considering forms of landscape connectivity relevant to each dispersal mode. Second, the landscape variables that emerged as important to each dispersal mode were generally consistent with our predictions based on putative dispersal vectors, and included interactive effects that allude to the altered efficacy of animal dispersal in invaded landscapes. Our results have broad implications for understanding how ecological communities reorganize as landscapes are fragmented, patches are lost, and the function of dispersal life histories is altered.

## Introduction

Ecologists have long-sought to quantify the importance of dispersal limitation in ecological communities (Borcard et al. 2004; Gilbert and Lechowicz 2004; Cottenie 2005) for two main reasons. First, the presence of dispersal limitation can cause local species richness to fall short of what a given environment can support (Germain et al. 2017), and second, the failure of propagules to reach suitable sites generates spatial turnover in species richness and composition that contribute to regional biodiversity (Hurtt and Pacala 1995; Mouquet and Loreau 2003). Towards this goal, numerous statistical tools have been developed to isolate the relative explanatory power of dispersal vs. environment from field data (*e.g.,* (Peres-Neto et al. 2006; Prugh 2009)), typically testing for and interpreting an effect of distance among local communities on species occupancy, richness, or composition as evidence of dispersal limitation (Hanski 1994*a*; Cottenie 2005; Prugh et al. 2008). If spatial distance among sites is assumed to be the best proxy of restricted dispersal, then the absence of significant distance effects on diversity patterns is interpreted as evidence that dispersal is not limiting at the spatial scales examined (*e.g.,* (Freestone and Inouye 2006)) – that is, that species have access to all habitat patches, and that variation in species occupancy and richness patterns reflect variation in local environmental conditions.

An alternative but rarely considered explanation for non-significant distance effects is that isolation by distance is not the spatial variable most relevant to dispersal – habitat patches might be close in space but poorly connected by dispersal due to other landscape features, such as physical barriers. Though this idea has been explored in aquatic ecosystems for which there is obvious network structure (*e.g.,* riverine networks (Beisner et al. 2006; Brown and Swan 2010)) or directionality to dispersal (*e.g.,* water currents (White et al. 2010)), it has not been explored in terrestrial systems for which dispersal barriers may be cryptic and thus difficult to identify and measure. Although isolation by distance is likely the most important factor limiting dispersal in many ecosystems (*e.g.,* oceanic islands), exploring alternative dispersal pathways can reveal hidden constraints to how species move across and interact with their landscapes, and might explain why distance effects are generally weak in terrestrial ecosystems; a recent synthesis of 1,015 animal surveys found that spatial isolation was a poor predictor of patch occupancy for most species (Prugh et al. 2008).

If dispersal is constrained in a greater range of ways other than by distances among habitat patches, then, intriguingly, species might differ in how they experience the spatial connectedness of the same physical landscape based on dispersal life histories (Beisner et al. 2006). In plants, for example, species possess a range of adaptations to disperse, called dispersal syndromes or “modes”, such as dispersal by gravity, animals, or wind. Previous research of understory herbs in aspen stands clearly demonstrates that dispersal mode dictates how constrained plant species distributions are by the size and spatial isolation of habitat patches, even without accounting for additional sources of trait variation (*e.g.,* seed size) among species within dispersal modes (Jones et al. 2015). In that study, however, the effect of dispersal mode on species distributions was not consistent with a simple difference in dispersal ability among modes (*i.e.,* dispersal ability: gravity < wind < animal (Jones et al. 2015)), as hypothesized if distances among habitat patches was the only cause of spatial isolation. We contend that linking species distributions to the spatial distribution of dispersal vector movement might be the missing piece needed to understand the mechanisms that underlie the spatial distribution and composition of biodiversity, for plants and potentially other terrestrial organisms. Identifying spatial constraints on species distributions is key to understanding the processes that underlie fundamental patterns in ecology, such as species-area relationships (Shen et al. 2009), as well as to forecast how ecological communities might reorganize as the spatial and environmental structure of landscapes is altered by humans (Gonzalez et al. 2011; Frishkoff et al. 2016).

We explored alternative forms of landscape connectivity to understand the distribution of biodiversity in a natural patch-network of plants that associate with serpentine soils. Serpentine soils form via the emergence and erosion of the Earth’s mantle into discrete patches embedded within a matrix of non-serpentine soil. Serpentine soils are hypothesized to act as “islands” of refuge for native plant species to escape the “sea” of European grasses that now dominate Californian landscapes (Harrison and Rajakaruna 2011; Gilbert and Levine 2013). The annual plant communities that associate with serpentine soils are an emerging model system to understand the mechanisms that underlie the spatial scaling of biodiversity (Anacker and Harrison 2012; Germain et al. 2017), the interaction between local and regional processes (Harrison 1999; Harrison et al. 2006), and the community impacts of species invasions (Gilbert and Levine 2013; Case et al. 2016). Recent experimental work has demonstrated that dispersal limits plant diversity at our study site (fig. 1), yet we find no evidence of spatial distance as a proxy for dispersal limitation through our observational data (table A2); this contradiction motivates our examination of other landscape features relevant to dispersal. Specifically, the absence of tall vegetation in serpentine grasslands allowed landscape features, such as hydrological networks and animal paths, to be captured via satellite imagery (fig. 2).

**Figure 1:**
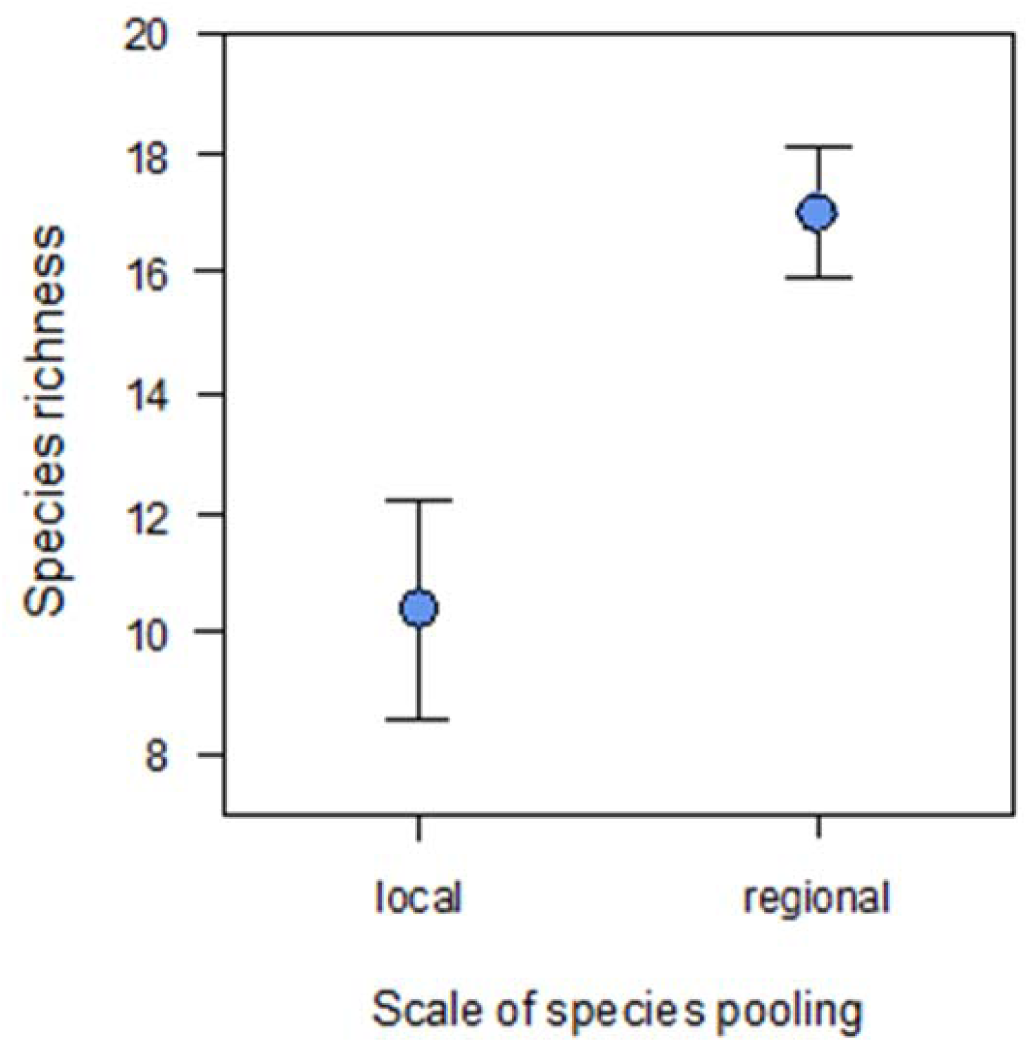
Experimental evidence of dispersal limitation via pooling seed banks within (local) or among (regional) habitat patches to enhance dispersal; on average, we see an increase of 7 species per habitat patch with regional pooling (*F_1,4_* = 15.2, *P* = 0.0175). The data presented here is subsetted from a larger dataset (Germain et al. 2017) to include only sites within the same region as our current survey, and only treatments that received locally-mixed species pools (5 m spatial scale) and those mixed among sites within the regional extent of our survey (100 m spatial scale).

**Figure 2:**
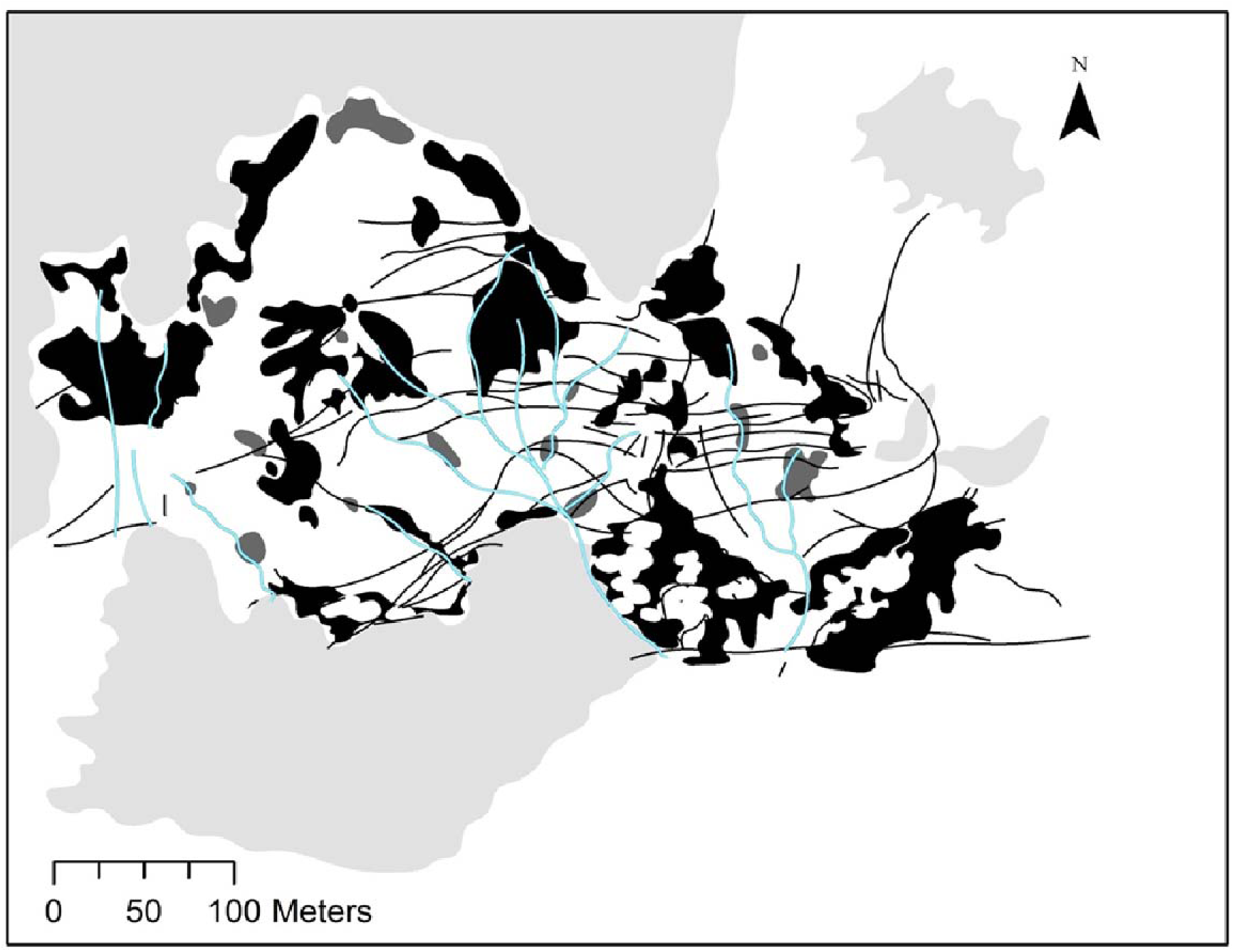
Map of sampled (black) and unsampled (dark gray) habitat patches at our 18-ha study site within McLaughlin Reserve, CA. Thin black lines are animal paths and thick blue lines are surface runoff, both traced from satellite imagery. The habitat matrix (white) was non-serpentine soils dominated by European grasses, such as *Avena barbata*, and the site boundaries were either road or chaparral (light grey).

We surveyed plant communities on serpentine patches and in the non-serpentine matrix, categorized species by dispersal mode, and estimated habitat patch characteristics relevant to different modes of dispersal. We used these data to answer two questions: (1) Are there landscape features that characterize habitat patch connectivity better than distance among habitat patches, and (2) does dispersal mode influence how species respond to these features? If species experience different landscape constraints, then we predict that the richness of species belonging to different dispersal modes will be highest in habitat patches highly connected by their respective dispersal vectors. Specifically, we predict that hydrological networks, animal paths, and distance would explain the richness of species dispersed by gravity, animals, and wind, respectively. As a case study, we also explore the spatial distribution of patch occupancy patterns of *Plantago erecta* (California plantain), a small-statured annual with seed morphologies consistent with a mixed dispersal strategy (*i.e.,* dispersal via water and animals (Germain et al. 2017)). If our models are correct, then we predict that *P. erecta*’s distributions would be explained by forms of habitat connectivity shown to be important to both dispersal modes.

Our analyses of species richness fall into a general class of ‘incidence function’ models (Prugh 2009), the basis of which was first developed by Levins (Levins 1969) and later adapted by Hanski (Hanski 1994*a*, 1994*b*) to test species’ extinction and colonization as a function of patch size and isolation by distance, respectively. These models have achieved broad success at understanding the population and metapopulation persistence of a diversity of organisms in fragmented landscapes (*e.g.,* butterflies [28], pikas (Moilanen et al. 1998)), with applications to landscape management and conservation planning (Wahlberg et al. 1996).

## Materials and Methods

### Study System

Our study took place at the 2800-ha McLaughlin Natural Reserve (http://nrs.ucdavis.edu/mcl/) in Northern California, at the boundary of Lake, Yolo, and Napa counties (38°51′47.01″N, 122°21′48.87″W). The landscape is characterized by patches of serpentine soil interspersed among a matrix of non-serpentine soil. Serpentine (ultramafic) soils are derived from the Earth’s mantle in regions where it becomes exposed, such as along the San Andreas Fault, and are identified by Ca/Mg ratios < 1 (Anacker 2014). Calcium is essential to plant growth, and is captured less efficiently in the presence of magnesium. Low Ca/Mg ratios, coupled with low soil fertility, high heavy metal content, and poor soil moisture retention, present a harsh growing environment for plants. Yet, serpentine soils support a rich diversity of native and endemic plant species (Anacker 2014), and are hypothesized to act as spatial refugia for native species to escape the competitive effects of the exotic European plants that now dominate the non-serpentine matrix (Gilbert and Levine 2013).

### Field Survey and Data Collection

Plant surveys and all fieldwork were conducted in early May 2017, at approximately peak flowering. We haphazardly selected 28 serpentine habitat patches out of all 42 patches in a 18 ha region of the reserve, ranging from 31 to 4533 m^2^ in size and 0.75 to 356 m away from their nearest neighbor patch (fig. 2). At each patch, we surveyed a transect of five 0.75 × 0.75 m^2^ plots: one plot in the patch centre, one plot halfway between each edge and the patch centre, one plot 1 m into the non-serpentine matrix, and one plot 5 m into the matrix (fig. A1). In other words, the distances among plots within patches were scaled by patch size, whereas the two matrix plots were fixed distances from the patch edge. We recorded the presences of all species in each plot, and made note of species that covered more than 25% of a plot by area (usually one to three species). In total, 77 plant species were present in our surveys, 72 of which could be identified; the five unidentified species occurred once each, had no distinguishing features to assess dispersal mode with certainty (*i.e.,* only a single basal leaf) and were discarded from all analyses that required information on dispersal mode. Sampling the same total area for all habitat patches regardless of patch size is a standard sampling method to prevent confounding patch size with sampling intensity (Cook et al. 2002).

Species’ dispersal modes (dispersal via wind, gravity, or animals) were categorized based on previous research (Spasojevic et al. 2014) and updated here based on seed/diaspore morphology and if more detailed information on dispersal modes was available (table A1). Wind-dispersed species were identified by the presence of a pappus or seed wings, whereas animal-dispersed species had morphologies for attachment to passing animals, such as burrs, awns, or hairs. Species categorized as gravity dispersed had seeds that lacked any apparent mechanism for dispersing by wind or animals, and tended to have smooth, spherical diaspores conducive to downslope dispersal via rain and gravity. We include ant-dispersed species as gravity dispersed given that ants disperse seeds at very small spatial scales and are unlikely to contribute to regional occupancy patterns (Thomson et al. 2011), as well as species with reduced pappi that were biomechanically unlikely to confer wind dispersal (*e.g., Lasthenia californica*). One species, *Plantago erecta* (California plantain), was previously categorized as being dispersed by water (Spasojevic et al. 2014). However, *P. erecta* seeds produce a sticky mucilage that might also allow dispersal by animals (observation noted in (Germain et al. 2017)). As such, we categorize this species as being animal-dispersed but also explore the occupancy patterns of this species in depth as a case study of a species with two potential dispersal modes.

Species were additionally categorized as patch- or matrix-associated (table A1) to identify and account for species that were unlikely constrained to serpentine habitat patches (Cook et al. 2002; Jones et al. 2015). ‘Matrix-associated’ species included both matrix specialists and generalists that show no affinity for habitat type. Species were considered matrix-associated if they were equally or more common in matrix plots than in the patch plots. Twelve species met these criteria, including *Avena fatua*, *Bromus hordeaceus*, and *Lotus wrangelianus*. Species richness (fig. A2*A*) and composition (fig. A2*B*) differed among serpentine habitat patches and the non-serpentine matrix (both *P* < 0.001), confirming that serpentine plant communities are distinct and thus constrained to the serpentine habitat-patch network.

We estimated habitat patch characteristics in the field and using ArcGIS on GoogleEarth images. In ArcGIS, we delineated all serpentine patches within our study region, including the 28 surveyed patches and 14 unsurveyed patches; these delineations allowed us to calculate patch size and patch connectivity. Patch connectivity was estimated using edge-to-edge distances between patch *i* and all other *j* patches (including the unsampled patches), weighted by a negative exponential dispersal kernel using eq. 1 (Hanski 1994*a*, 1994*b*; Jones et al. 2015):

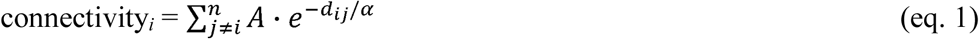

where *A* is the area of patch *j*, *d* is the Euclidean distance in meters between patch *i* and *j*, and *α* is the mean dispersal distance, set to 5 m for all species. In other words, patch *i* is most connected when it is in close proximity to many large patches. Our connectivity measure falls into a general class of measures called incidence function models, which have been shown to perform equally well or better than alternative measures (*i.e.,* nearest-neighbour or buffer measures (Prugh 2009).

We estimated two alternative measures of connectivity that we hypothesized could be more appropriate for plant species that are dispersed by animals or gravity. For species that are dispersed by animals, we traced deer trails that were observable via GoogleEarth (fig. 2) using ArcGIS, and used the number of trails that intersected habitat patches to estimate patch connectivity via animals. Deer exhibit path fidelity, following the path of least resistance, which is especially true in topographically challenging landscapes such as our study area, and create “highways” for a diversity of other animals to traverse landscapes, including rabbits and turkeys (Sindorf 2009). For plant species that are dispersed by gravity/water, we estimated hydrological connectivity by tracing the network of surface streams, and for each habitat patch, summed the area of all *j* habitat patches upslope from and connected via surface streams to each patch *i*. Elevation of habitat patches ranged 450-550 m a.s.l., small enough for elevational clines in climate to be unimportant. Although in some systems, habitat patches at the base of an elevational cline have increased resource inputs and thus higher productivity, productivity was not correlated with elevation (*slope* < 0.01, *P* = 0.465) or hydrological connectivity (*slope* = −0.03, *P* = 0.458) in our dataset. We estimated productivity as a composite measure [(1 - proportion of bare ground) × vegetation height] to non-destructively estimate the volume of plant material in each plot.

### Statistical Analyses

To test if species composition in serpentine habitat patches was distinct from the surrounding non-serpentine matrix, we used linear mixed effects models to test differences in species composition among plots in serpentine habitat patches (“patch plots”), 1 m into the habitat matrix (“edge plots”), and 5 m into the habitat matrix (“matrix plots”). To do so, we first performed a principal coordinates analysis using Jaccard’s distances on the plot-level presence/absence data. The first and second axis scores were used as response variables in separate analyses with fixed effects of habitat type (*i.e.,* patch, edge, matrix) and ‘patch id’ included as a random effect to account for the non-independence of the five plots transecting each habitat patch. The ‘glht’ function in R package ‘multcomp’ was used to to perform a Tukey’s tests of compositional differences among all pairwise treatment combinations (patch vs. edge, patch vs. matrix, edge vs. matrix).

To examine the relationship between species richness and patch connectivity, we performed a generalized linear mixed effects model with species richness as the response, fixed effects of dispersal mode, connectivity by distance, connectivity by animals, connectivity by hydrology, patch size, and all interactions, and random effect of ‘patch id’ to account for non-independence of replicate plots within a single patch. Because there were significant higher-order interactions between dispersal mode and all connectivity measures (table A2), we performed separate analyses of biogeographic predictors for each dispersal mode. This type of analysis is highly prone to type I errors (poisson-distributed data with interacting continuous predictors), so we took several steps to identify reduced models that best fit the data. First, we performed backwards selection on each full model using the ‘step’ function in the ‘stats’ package; ‘step’ sequentially drops higher order interactions until the reduced model that yields the greatest model fit (lowest AIC score) is attained. However, ‘step’ can arrive at a local minimum in AIC score that does not reflect the global minimum, which was likely for our data given the presence of significant but biologically implausible four-way interactions. For this reason, we applied ‘drop1’ to the ‘step’-reduced model to identify variables that did not significantly improve model fit even if their inclusion led to a marginal decrease in AIC scores. We cycled between ‘drop1’ and ‘step’ until a model was obtained for which all variables significantly improved model fit and led to the lowest AIC scores out of all possible reduced models. We then used the function ‘Anova’ in the R package ‘car’ to test whether the reduced model was a significantly better fit to the data than an intercept-only model.

## Results and Discussion

Despite the emphasis that contemporary ecology places on dispersal as central to the dynamics and distribution of species in ecological communities (Hanski 1994*a*; Leibold et al. 2004), current empirical assessments of its role may not encompass the diversity of ways in which organisms experience landscape connectivity. In a patchy terrestrial plant community, we found no evidence of dispersal limitation using a distance-based estimate of patch connectivity, even after discounting the presences of species associated with the habitat matrix (*i.e.,* a non-significant effect of connectivity by distance; table A2), despite experimental evidence of its pervasiveness (fig. 1 with data from (Germain et al. 2017)). However, when separated species richness by dispersal mode (gravity, wind, animal), the spatial distributions of species richness generally corresponded to spatial patterns of dispersal vectors which connect habitat patches. We discuss these general findings, as well as several unexpected contingencies that provide a richer understanding of interacting dispersal vectors in serpentine grasslands and their altered efficacy in invaded landscapes.

Consistent with our prediction that the richness of gravity-dispersed species would be highest in patches highly connected by hydrology, hydrology was the only form of connectivity retained as a predictor after model selection for this group. However, the effect of hydrological connectivity was not simply a main effect, but rather, an interactive effect with patch size (*i.e.,* significant hydrological connectivity × patch size effect [*X^2^* = 6.37, *P* = 0.012]), such that species richness increased with each predictor only at low values of the other (*i.e.,* fig. 3*A*, steep slopes connecting points 1 to 2 and points 1 to 4, but shallow slopes connecting points 3 to 4 and points 2 to 3). Although we did not predict this interaction *a priori*, it suggests that large, hydrologically-connected patches are locally saturated (*i.e.,* response surface decelerates from points 1 to 3, note log-scale of axes) and that these two predictors act as compensatory pathways towards reaching saturation. Our findings are consistent with recent experimental work showing that dispersal only increases species richness in small habitat patches (Schuler et al. 2017), given that populations in small patches are more prone to stochastic extinctions (Gilbert and Levine 2017) which can be overcome via dispersal.

**Figure 3:**
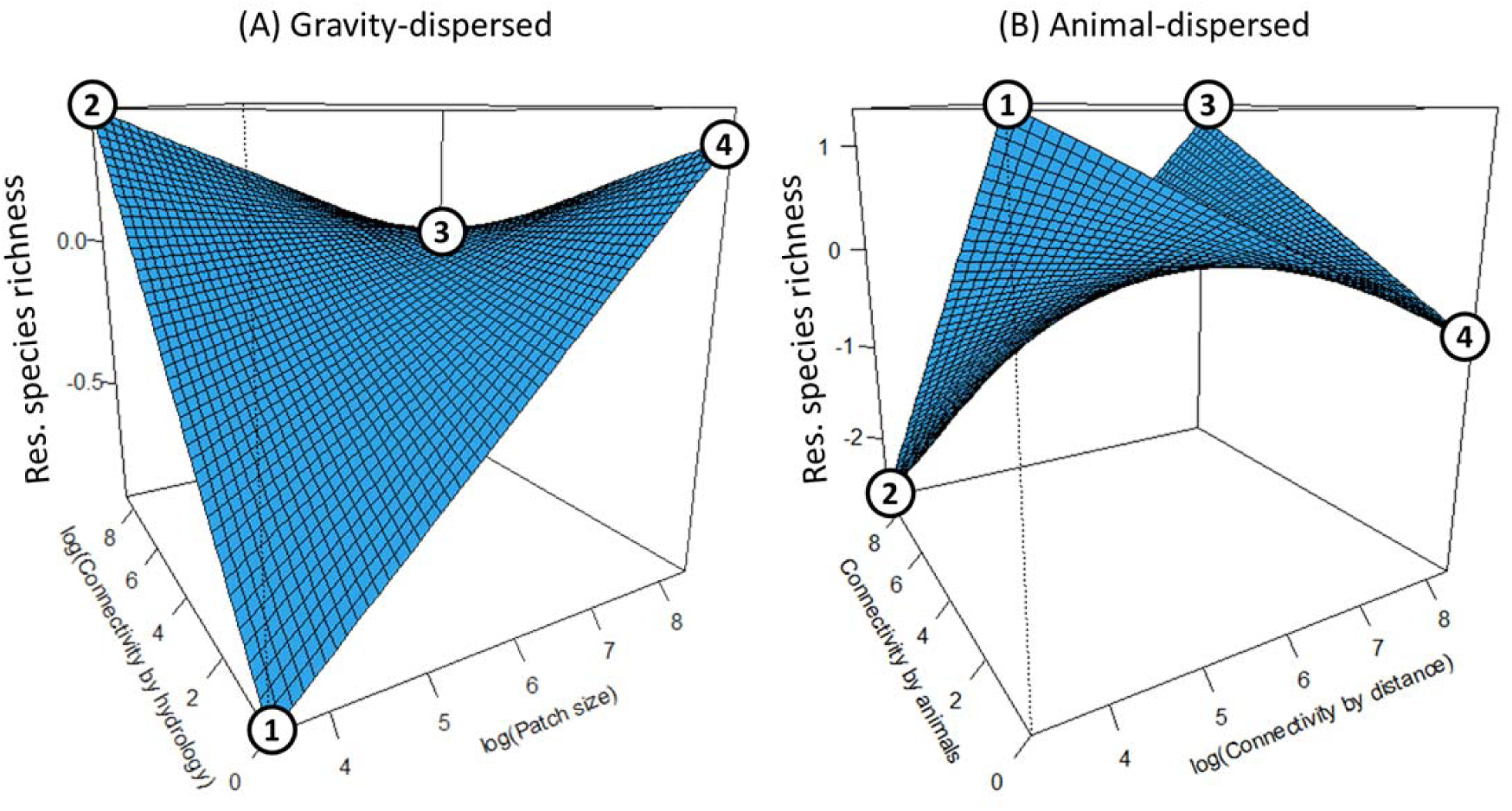
Species richness of (*A*) gravity-dispersed species, as a function of patch size and hydrological connectivity, and of (*B*) animal-dispersed species, as a function of connectivity by distance and animal connectivity. Numbered points connect different slopes to aid our description of the response surface in main text. Fitted response surfaces of species richness are shown for simplicity, after partialling out fixed effect of patch productivity and random effect of site ID, but residuals are shown in fig. A3.

The model that best fit the richness data of animal-dispersed species was one that, as predicted, included the appropriate vector of dispersal: connectivity by animals. However, as with gravity-dispersed species, the best-fit model also included an interaction, in this case between connectivity by animals and connectivity by distance (*X^2^* = 7.06, *P* = 0.007), generating a complex response surface (fig. 3*B*). More species were found in serpentine patches intersected by many animal paths, but only when patches were in close proximity to one another (slope connecting points 3 to 4 in fig. 3*B*) – when patches were isolated, however, animals had a strongly negative effect (slope connecting points 1 to 2). What is driving the negative effect of animals in isolated patches? The answer is not likely herbivory, given that the animal-dispersed species in our dataset are generally tolerant of or well-defended against herbivory (*e.g.,* grasses, star thistle (table A1)) and given that connectivity by animals did not predict the richness of wind-dispersed species, a highly palatable group (*e.g.,* wild lettuce, dandelion (table A1)). Rather, we contend that the answer has more to do with the efficacy of animals as dispersal vectors in invaded landscapes. Seeds removed by animals in isolated habitat patches have a low probability of (i) being deposited in other habitat patches, compared to the inhospitable matrix, and of (ii) being rescued from extinction via dispersal from other patches; in other words, seeds are removed but not replaced. Additionally, many of the most noxious invaders in serpentine grasslands are animal dispersed (*e.g.,* common wild oat [*Avena fatua*], barbed goatgrass [*Aegilops triuncialis*]), adding the potential for invader impacts via animal-mediated dispersal from the non-serpentine matrix, even if those invaders exist only as sink populations (Schreiber and Kelton 2005). We argue that the negative effect of dispersal via animals is likely a contemporary phenomenon, given that, prior to invasion by European grasses, a greater proportion of the landscape was suitable to species that are now restricted to occur only on serpentine patches (Gram et al. 2004; Gilbert and Levine 2013). Further support for this hypothesis comes from our finding that species richness increases with connectivity by distance only in patches highly connected by animals (slope connecting points 2 and 3), reinforcing animals as dispersal vectors, connecting patches that would otherwise be unconnected reduce close proximity.

Although we predicted that the richness of wind-dispersed species would increase with connectivity by distance, given that wind moves in all directions in topographically complex landscapes (McNider and Pielke 1984), such as our study site, we instead found that an intercept-only model best fit the data. This finding has three possible explanations, the first being that wind-dispersed species are simply not dispersal limited at the spatial scale of our surveys, and the second being the possibility that we have not adequately captured spatial variation in the movement of seeds by wind. Although we cannot weigh these two alternate explanations against each other, what we can say is that there is a high degree of variation in species richness and composition among patches for this dispersal group, including some patches that lack species from this group altogether. High spatial turnover (β diversity) without evidence of dispersal limitation implicates the role of local processes (Germain et al. 2013), such as environment, herbivory, competition, or stochasticity. However, a more detailed examination of dispersal kernels and constraints for this group are needed.

The third explanation is that trait differences among wind-dispersed species, for example, short vs. tall species (Thomson et al. 2011), caused additional variation in how species experience landscape connectivity. Testing this possibility would require separate analyses of species occupancy patterns for multiple species; our data is not amenable to such an analysis, because only two wind-dispersed species occupied enough patches to reasonably fit an incidence function model (MacKenzie et al. 2005). Coarsely, though, the most common wind-dispersed species (*Microseris douglasii*), observed in 22 of the 28 sampled patches, was average in terms of plant height and the ratio of seed size to dispersal structure, though did have the largest seeds (fig. A4). Large-seeded wind-dispersed plant species disperse farther on average (Thomson et al. 2011), thus seed size differences may contribute to regional occupancy patterns for this group.

As predicted, the distribution of *P. erecta*, a common small-statured annual with seed morphologies consistent with a mixed dispersal strategy (*i.e.,* dispersal via water and animals (Germain et al. 2017)), was explained by patch characteristics consistent with both dispersal modes. Specifically, occupancy patterns of this species were influenced by a three-way interaction between connectivity by distance, hydrology, and animals (*X* ^2^ = 5.70, *P* = 0.017), as well as positive main effects of hydrology and animals (both *P* ≤ 0.001; table 1). When patches were well-connected by hydrology, the response surface of the probability that *P. erecta* was present in patches resembled that of richness of animal-dispersed species (fig. A5*B* vs. fig. 3*B*). However, when patches were poorly connected by hydrology, occurrence probabilities generally increased with connectivity by animals (fig. A5*A*). This in-depth examination of single-species occupancy patterns demonstrates consilience among approaches, where connectivity measures identified as important to different dispersal modes in the community level data also emerge as important predictors of a species with a mixed dispersal strategy.

**Table 1:** Occupancy of *Plantago erecta*, a species with a mixed dispersal strategy, is explained by connectivity measures intermediate to those exhibited by species with animal and gravity dispersal

| Variables | Slope | $X^2$ value | *P*-value |
| --- | --- | --- | --- |
| size | -2.15 | 0.06 | 0.812 |
| distance | 13.30 | 0.31 | 0.578 |
| animals | 30.99 | 10.44 | **0.001** |
| hydrology | 13.64 | 36.29 | **<0.001** |
| distance × animals | -4.70 | 2.15 | 0.143 |
| distance × hydrology | -2.53 | 1.83 | 0.176 |
| size × hydrology | 0.47 | 2.82 | 0.093 |
| animals × hydrology | -5.57 | 6.34 | **0.012** |
| distance × animal × hydrology | 0.85 | 5.70 | **0.017** |

*Note:* Significant *P*-values are in bold typeface. This reduced model provided a significantly better fit to the occupancy data than an intercept-only model, despite requiring an additional 9 degrees of freedom (model comparison: $P = 0.012$ ).

Habitat fragmentation is the primary driver of biodiversity loss worldwide (Crooks et al. 2011). In serpentine plant communities and many other ecosystems, fragmentation has occurred via the widespread invasion of non-native species, with native species now relegated to small isolated “refuge” habitat patches. Though species in refuge patches may be safe from direct competition with invaders, diversity is still challenged with the indirect effects of reduced colonization (Gilbert and Levine 2013). The extreme harshness of the competitive effect in the non-serpentine matrix is clear if we consider that (i) plots in the non-serpentine matrix were 7.1 times more productive than serpentine plots yet contained 2.2 fewer species on average (fig. A2*A*), and that (ii) there was no difference in species composition among plots 1 m vs. 5 m into the matrix (grey vs. white points in fig. A2*B*) even though 1 m is within the dispersal capacities of most species. In order to prevent the non-random loss of some species over others (*e.g.,* plants dispersed by animals), landscape management plans may need to consider alternate multiple forms of habitat connectivity. Californian landscapes were invaded ~200 years ago, meaning that current communities may already reflect the compositional reorganization of some groups over others, a hypothesis that can be tested experimentally.

## Conclusion

Characterizing habitat connectivity is fundamental to understanding how dispersal contributes to biodiversity patterns (Leibold et al. 2004), as well as to landscape planning for conservation (Crooks and Sanjayan 2006). In a serpentine grassland, we uncover cryptic dispersal networks by linking species’ dispersal life histories to dispersal vector movement. Our results suggest that ecologists should more carefully consider whether the absence of significant distance effects truly represents an absence of dispersal limitation *vs.* a failure to capture landscape variables that are most limiting to dispersal. Additionally, our finding that animal dispersal reduced diversity in isolated habitat patches points towards the altered functioning of ecological networks in invaded landscapes. Real landscapes include complex spatial flows of energy and matter, which as we demonstrate, sets up ecological opportunity for organisms to differ in how they interact with and experience the same landscape.

## Acknowledgements

We thank Diane Srivastava and the Srivastava lab for comments on the manuscript, as well as Roi Holzman for references on connectivity in the marine literature. Research funding was provided to R.M.G. by the Biodiversity Research Centre at UBC and the Killam Trust.

## Author’s Contributions

RMG conceived of the study, RMG, NTJ, TNG designed the sampling design and performed fieldwork, NTJ did the GIS, RMG and NTJ analyzed the data, RMG, NTJ, TNG wrote the paper.

